# Nanopore adaptive sampling effectively enriches bacterial plasmids

**DOI:** 10.1101/2022.10.03.510741

**Authors:** Jens-Uwe Ulrich, Lennard Epping, Tanja Pilz, Birgit Walther, Kerstin Stingl, Torsten Semmler, Bernhard Y. Renard

## Abstract

Bacterial plasmids play a major role in the spread of antibiotic resistance genes. However, their characterization via DNA sequencing suffers from the low abundance of plasmid DNA in those samples. Although sample preparation methods can enrich the proportion of plasmid DNA before sequencing, these methods are expensive and laborious, and they might introduce a bias by enriching only for specific plasmid DNA sequences. Nanopore adaptive sampling could overcome these issues by rejecting uninteresting DNA molecules during the sequencing process. In this study, we assess the application of adaptive sampling for the enrichment of low-abundant plasmids in known bacterial isolates using two different adaptive sampling tools. We show that a significant enrichment can be achieved even on expired flow cells. By applying adaptive sampling, we also improve the quality of *de novo* plasmid assemblies and reduce the sequencing time. However, our experiments also highlight issues with adaptive sampling if target and non-target sequences span similar regions.

## Introduction

Infectious diseases caused by bacterial pathogens have lost their threat to people living in high-income countries due to the discovery of antibiotic drugs within the last 70 years. However, adaptation processes within bacteria cause these drugs to lose their effectiveness in treating infectious diseases. The emergence of such antimicrobial resistance (AMR) already poses a significant threat to public health, with an estimated 4.95 million deaths associated with bacterial AMR in 2019 (1), and will even worsen, with around 10 million expected deaths per year by 2050 (2, 3). Besides vertically passing antimicrobial resistance genes (ARG) to their offspring, bacteria can also transfer ARGs across the bacterial population by horizontal gene transfer. This process is mediated via mobile genetic elements (MGE), such as plasmids, which are epichromosomal DNA elements unique to bacteria (4, 5). Plasmids are a major driver in the spread of ARGs in bacterial populations (6) and have recently been found to accelerate bacterial evolution by enhancing the adaptation of the bacterial chromosome (7).

Classifying plasmid types is crucial to understanding antibiotic resistance transmission between bacteria. Several recent studies have shown the benefit of whole genome sequencing for classifying plasmid types (8, 9). In particular, the emergence of long-read sequencing by Oxford Nanopore Technologies (ONT) promises improvements for outbreak investigations due to its lower capital investment and shorter turnaround times (10, 11). However, these methods suffer from the small proportion of plasmid DNA within the sequenced samples, primarily if bacteria can not be cultivated in the lab (12). Therefore, a large proportion of the plasmids in such samples are probably missed, or the sequencing depth is insufficient to assemble them correctly (13). Thus, additional sample preparation steps are required to isolate or enrich plasmids before DNA sequencing, but they are too expensive and laborious for applications in clinical diagnostic settings.

While nanopore sequencing has been shown to reconstruct plasmids accurately (14), the technology offers a feature called adaptive sampling (AS) that has the potential to improve plasmid classification. First described in 2016 by Loose et al., nanopore adaptive sampling has been increasingly used for *in-silico* target enrichment within the last two years. Here, DNA molecules can be rejected from individual nanopores if the corresponding sequence is not interesting for downstream analysis. Pulling out unwanted DNA frees the nanopore for the following molecule to be sequenced and reduces the time spent sequencing uninteresting DNA fragments. Different tools implement adaptive sampling (16–18), using dynamic time warping (UNCALLED), read mapping (Readfish, MinKNOW) or k-mer-based (ReadBouncer) strategies, all performing rejection decisions by analyzing the first 160 to 450 base pairs of each read. Recently, deep-learning-based tools like SquiggleNet and DeepSelectNet have also been developed, addressing host depletion in human microbiome samples (19, 20). The potential enrichment reached by using adaptive sampling was already shown, and even mathematical models that predict the enrichment factor were recently described by some groups (17, 21, 22). In one study, Marquet et al. could enrich the microbiome in human vaginal samples by depleting host DNA. Further, Kipp et al. used adaptive sampling to enrich bacterial pathogens in tick samples, while Viehweger et al. even enriched single ARGs in human microbiome samples.

In the present study, we investigate the efficiency of adaptive sampling to enrich plasmid sequences in five different bacterial isolates. For this purpose, we used two adaptive sampling tools, which were shown to have high read classification performance, namely the built-in adaptive sampling feature of MinKNOW (referred to as ‘MinKNOW’ in this manuscript) and ReadBouncer (18, 19). Both tools use a combination of base-calling with ONT’s `Guppy` and read classification on sequence level. While MinKNOW’s adaptive sampling feature is based on the Readfish (17) scripts and uses minimap2 to map read prefixes against a given reference sequence set, ReadBouncer utilizes pseudo-mapping based on k-mers and interleaved Bloom Filters for making rejection decisions. We refrained from using UNCALLED because Bao et al. showed that the combination of base-calling and mapping has a higher read classification accuracy than UNCALLED. In order to increase sustainability and reduce sequencing costs, we also investigate whether enrichment of plasmids can be achieved with adaptive sampling on expired flow cells with reduced active pores. Finally, we evaluate the effective plasmid enrichment by comparing it to the predicted enrichment calculated by the mathematical model proposed by Martin et al. and demonstrate the usefulness of adaptive sampling for plasmid assemblies.

## Methods

## Methods

### Culture and DNA extraction

*Campylobacter* strains were streaked on Columbia Blood agar (Oxoid, Thermofisher Scientific, USA) and incubated at 42 °C under microaerobic atmosphere. *Enterobacter, Salmonella* and *Klebsiella* strains used in this study were streaked out on Luria Bertani (LB) plate and incubated over night at 37 °C. DNA extraction for *Campylobacter jejuni* (GCF_008386335.1) was done using the MagAttract HMW Genomic Extraction Kit (Qiagen). For *Salmonella enterica* (GCA_025839605.1), *Campylobacter coli* (GCF_025908295.1) (25), *Klebsiella pneumoniae* (GCF_025837075.1) and *Enterobacter hormaechei* (GCF_001729785.1) DNA was extracted using the QIAamp DNA Mini Kit (Qiagen, Hilden, Germany) according to the manufacturer’s instruction. The total amount of DNA was quantified using a Qubit fluorometer (Thermo Fisher Scientific) and frozen at -80ºC until further analysis.

### Library preparation and sequencing

Sample preparation was performed according to the manufacturer’s instructions without any optional pre-enrichment steps or size selection using the Rapid Barcoding Kit SQK-RBK004. Different barcodes were used for each of the bacterial isolate samples to correctly assign sequenced reads in the data analysis. Since we used expired flow cells with less expected overall sequencing yield, we decided to only sequence two or three bacterial isolates on one flow cell. Finally, the barcoded samples were sequenced on an Oxford Nanopore MinION (Oxford, UK) using FLO-MIN106D(R9.4.1) flow cells. All sequencing experiments were started via ONT’s MinKNOW control software (version 4.5.0).

### *In-silico* enrichment via adpative sampling

We performed four sequencing runs using MinKNOW software v4.5.0 on an Nvidia Jetson AGX Xavier (512-core NVIDIA Volta GPU, 32GB LPDDR4X Memory) for 24 hours. In all experiments, we compared adaptive sampling with standard sequencing by dividing the flow cells into two parts: Adaptive sampling was performed on the first 256 channels, and standard sequencing was performed on channels 257 to 512. We used a new flow cell with 1153 active pores for the first run (ReadBouncer1) and sequenced two *Campylobacter* isolates using barcodes RBK01 and RBK02. For the second run (ReadBouncer2), we used an expired flow cell with only 636 active pores for sequencing the three barcoded bacterial isolates (*Enterobacter, Salmonella, Klebsiella*) using barcodes RBK03, RBK04 and RBK05. The third run (MinKNOW1) used the same *Campylobacter* samples as the first, but we performed sequencing on an expired flow cell with only 557 active pores. For the fourth run (MinKNOW2), we used the identical three bacterial isolates as for the second run and performed sequencing on an expired flow cell with only 718 active pores after the initial flow cell check.

On the first two flow cells, we performed adaptive sampling with `ReadBouncer` (18) using the chromosomal references of the bacterial isolates as depletion targets. Here, a k-mer size of 15, a chunk length of 250 bp, a fragment size of 200,000 bp, and an expected error rate of 5% were used as parameters for the read classification. `ReadBouncer`’s k-mer size parameter was chosen accordingly to the default k-mer size used for mapping with `minimap2` (26), which is used by `MinKNOW`’s adaptive sampling feature. The expected error rate reflects the current average per-read accuracy by ONT’s `Guppy` basecaller. The other two parameters are default parameters. For flow cells three and four, `MinKNOW`’s adaptive sampling feature was used, which is based on the `Readfish` (17) scripts and uses `minimap2` (26) to map read prefixes against a given reference sequence set for read classification. We built a `minimap2` index file (parameter `-x map-ont`) for these experiments, including the chromosomal reference sequences, which we used as depletion targets for adaptive sampling. Read prefixes classified as “chromosomal” were rejected from the pore, and decisions were written to log files by both tools, `ReadBouncer` and `MinKNOW`. In all experiments, `ReadBouncer` and `MinKNOW` used `Guppy` GPU basecaller (fast model, v6.0.6. Oxford Nanopore Technologies) for real-time base calling of the raw signal data received from the device after at most 0.4 seconds of sequencing.

## Data Analysis

All data analysis scripts were written in Python and R, and are freely available in the GitHub repository https://github.com/JensUweUlrich/PlasmidEnrichmentScripts.

All plots were created in R using ggplot2.

After the sequencing runs were finished, we basecalled and demultiplexed all raw data with `Guppy` GPU basecaller (super accuracy model, v6.0.6. Oxford Nanopore Technologies). `Guppy` trimmed barcodes and adapter sequences from the resulting nanopore reads during that process. Afterward, we computed read length metrics (see Table 1 and Figure 1) and created contour plots (see Figures 4 & 3) using the sequencing_summary files provided with the `MinKNOW` and `Guppy` output directories. Next, we mapped all demultiplexed and base-called reads against the reference genomes (including plasmid sequences) of the five bacterial strains using `minimap2 v2.19` (26) with parameter `-x map-ont`. Based on the mapping results, we could assign each mapped read to either the bacterial chromosome or plasmid(s) of one of the bacterial isolates to create Figure 2. We also used the mapping results to calculate the percentage of sequenced plasmid and chromosome base pairs after 24 hours for each bacterial sample, resulting in Figure 5. We further used the sequencing summary file to separate the reads by their species of origin and partitioned them to comprise the cumulative data from the beginning of each experiment up to 24 hours, separated by 30 minutes of sequencing, which resulted in 48 individual timepoint data sets. With this information, we calculated for each experiment time point *t* the plasmid enrichment by yield for each bacterial strain,

**Table 1.** Overview of read length metrics of the five bacterial isolates. The metrics were computed from reads sequenced on the control side of the flow cells where no adaptive sampling was applied.

| Reference | Flow cell ID | Mean Length | Median Length | Std. Deviation |
| --- | --- | --- | --- | --- |
| *Campylobacter jejuni* | ReadBouncer1 | 16,525.65 | 10,950 | 17,083.76 |
|  | MinKNOW1 | 13,360.65 | 8,192 | 15,072.50 |
| *Campylobacter coli* | ReadBouncer1 | 4,375.37 | 1,580 | 6,952.86 |
|  | MinKNOW1 | 5,536.69 | 2,736 | 6,894.11 |
| *Salmonella enterica* | ReadBouncer2 | 9,304.79 | 6,455 | 9,131.90 |
|  | MinKNOW2 | 8,679.22 | 5,952 | 8,629.98 |
| *Enterobacter hormaechei* | ReadBouncer2 | 8,909.21 | 6,239 | 8,845.49 |
|  | MinKNOW2 | 8,309.64 | 8,861 | 8,192.69 |
| *Klebsiella pneumoniae* | ReadBouncer2 | 9,379.93 | 6,512 | 9,180.22 |
|  | MinKNOW2 | 8,842.23 | 5,963 | 8,928.63 |

**Fig. 1.**
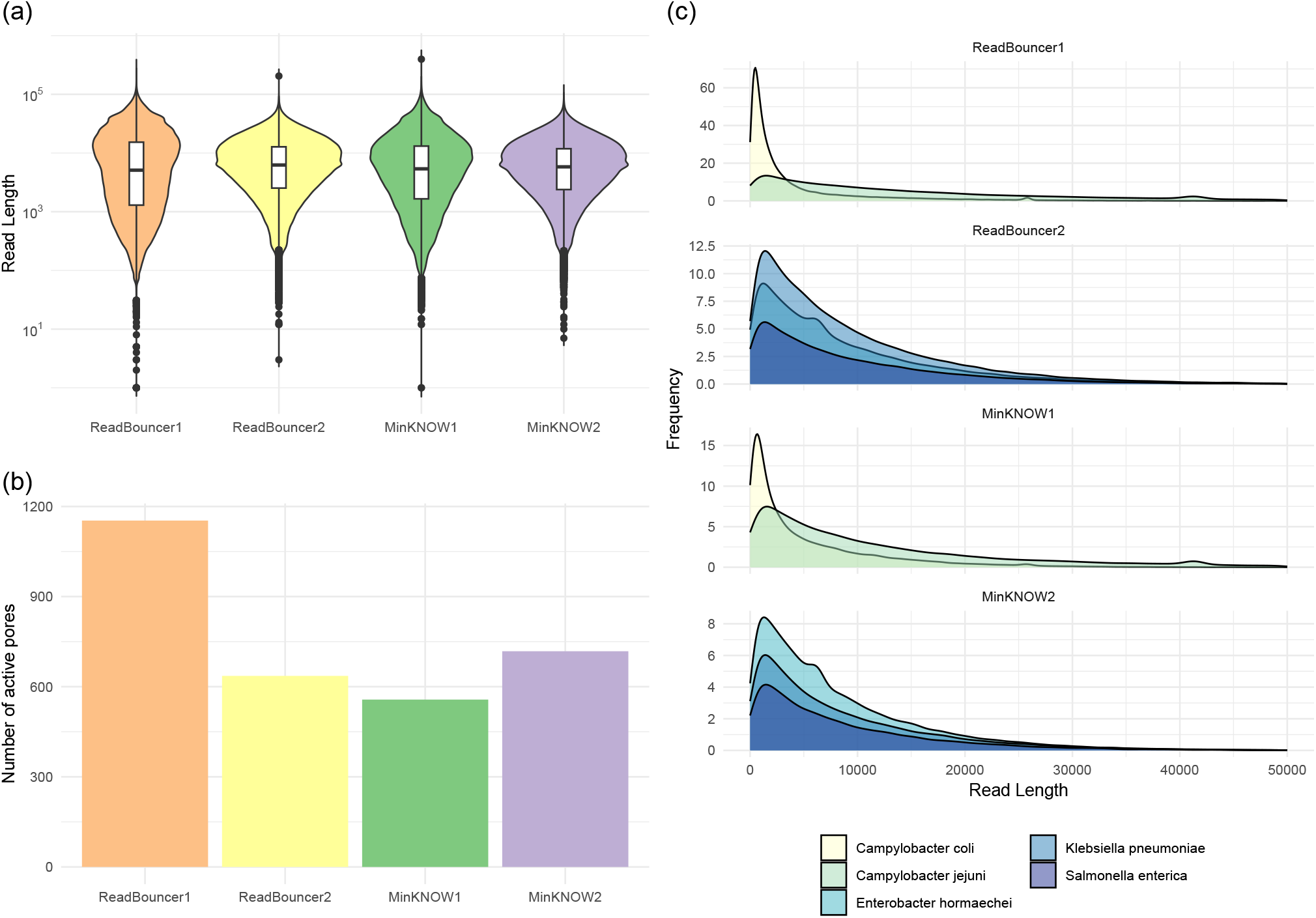
Evaluation of control regions from the first four sequencing runs. **(a)** Violin plots (log scale) of read length distributions. Box plots for read lengths are included within the violin plots. **(b)** Active pore count measured at the start of each sequencing run. ReadBouncer1 has the highest number of active pores because it was the only flow cell that was not expired. **(c)** Read length distributions for each species from control regions of the four sequencing runs.

**Fig. 2.**
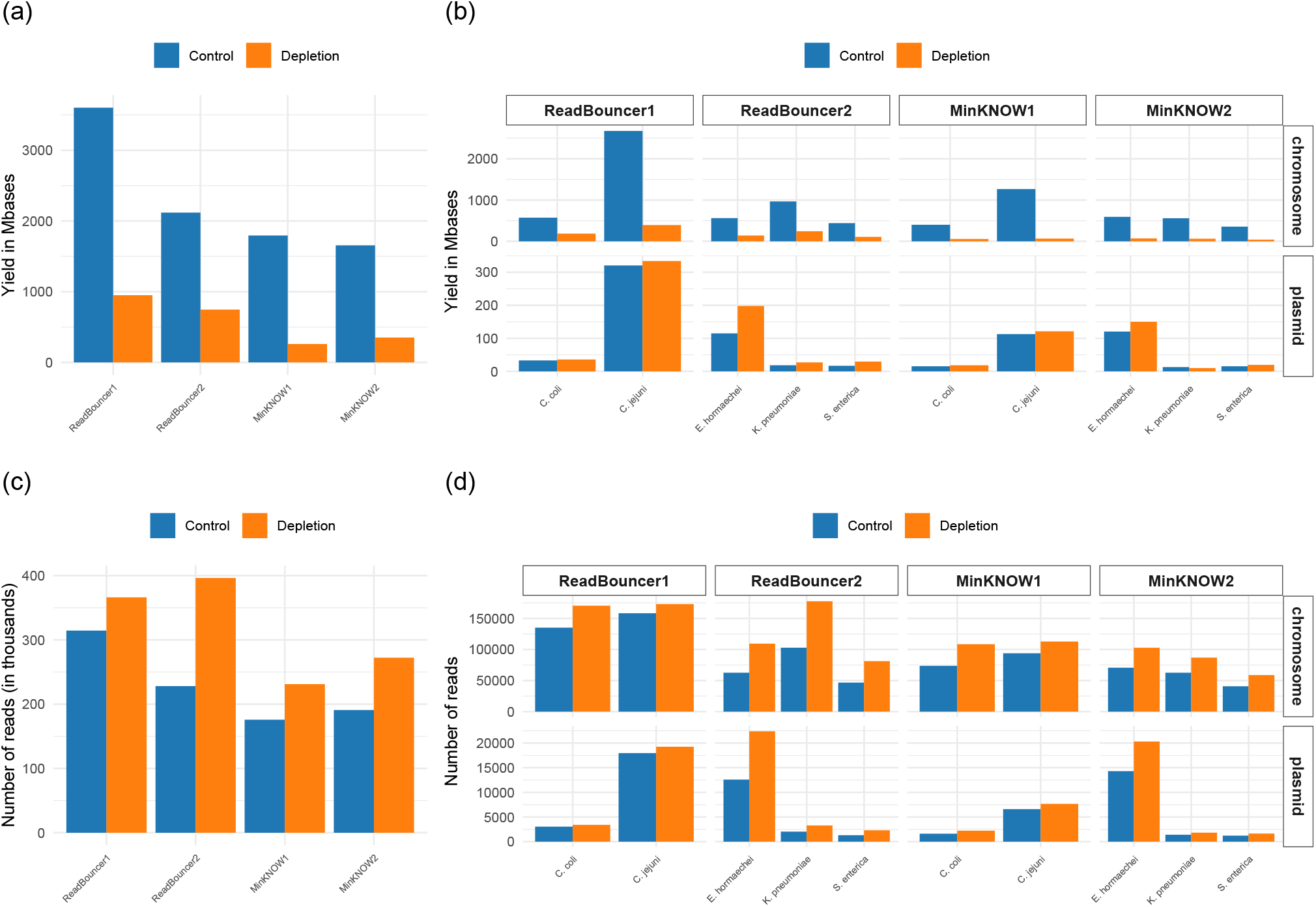
Comparison of flow cell yield in terms of sequenced base pairs and reads after 24 hours. **(a)** Yield in Megabases for each flow cell divided by control and adaptive sampling region (Depletion). **(b)** Yield in Megabases for each flow cell region divided by plasmid and chromosome. **(c)** Number of sequenced reads for each flow cell divided by control and adaptive sampling region (Depletion). **(d)** Number of sequenced reads for each flow cell region divided by plasmid and chromosome.

**Fig. 3.**
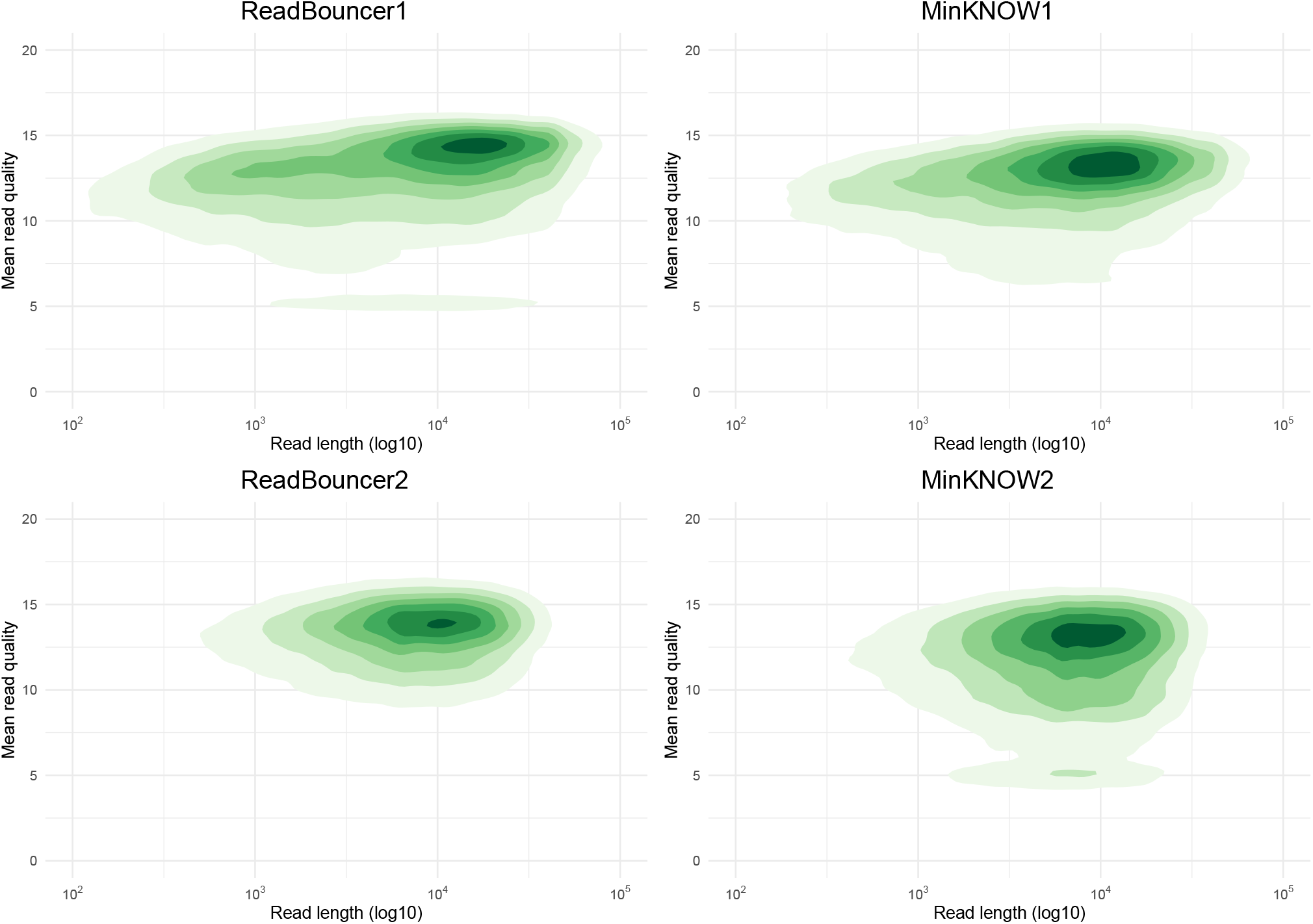
Contour plots of read lengths (log scale) against mean read quality for control regions of the four sequencing runs. Darker regions indicate a higher proportion of reads that fall into that slice. For example, ReadBouncer1 and MinKNOW1 have a higher proportion of reads with length above 10,000 base pairs than for ReadBouncer2 and MinKNOW2.

**Fig. 4.**
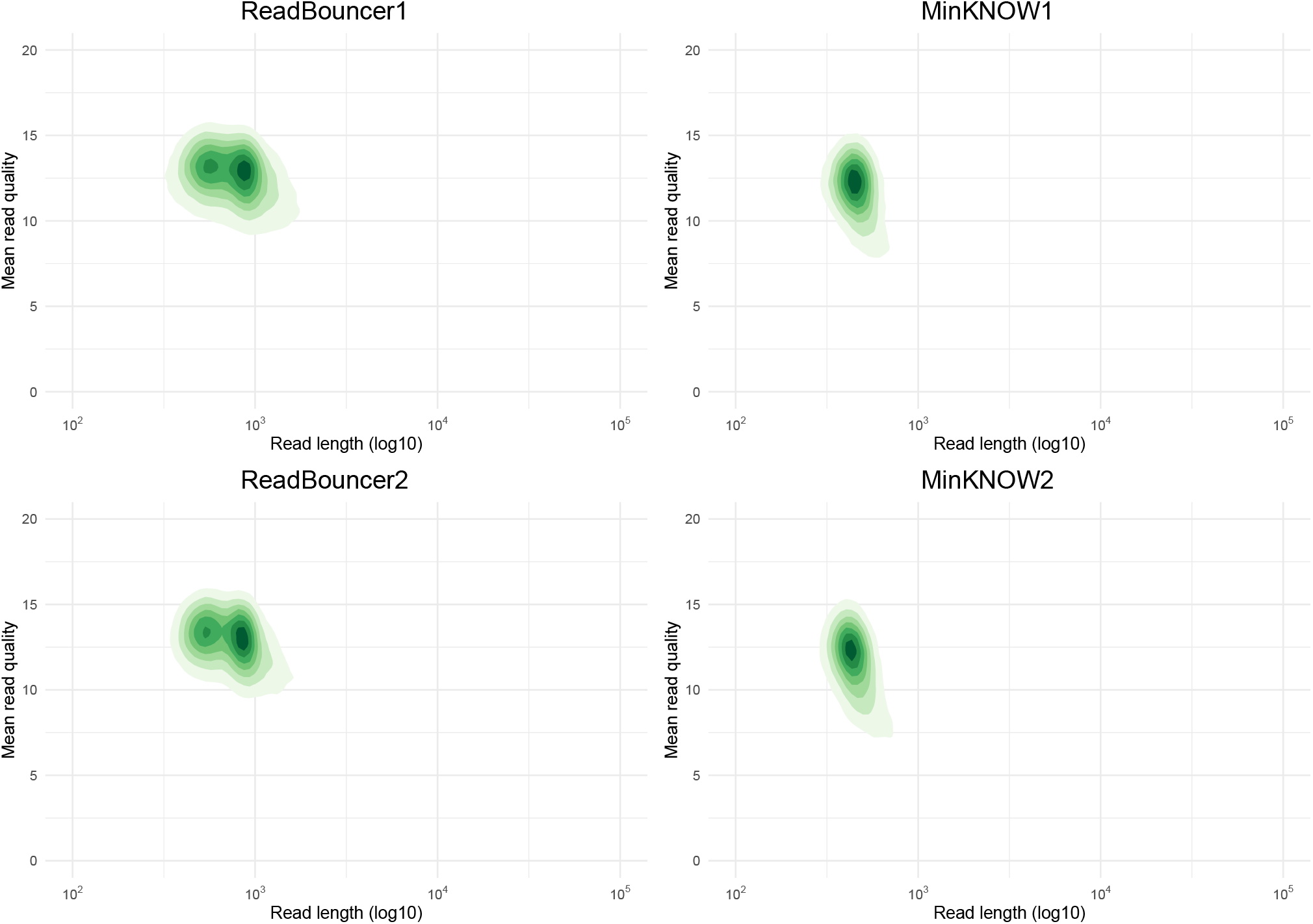
Contour plots of read lengths (log scale) against mean read quality for adaptive sampling regions of the four sequencing runs. Darker regions indicate a higher proportion of reads that fall into that slice. For example, most reads from sequencing runs MinKNOW1 and MinKNOW2 have read lengths of about 650 base pairs and a phred quality value of around 12.

**Fig. 5.**
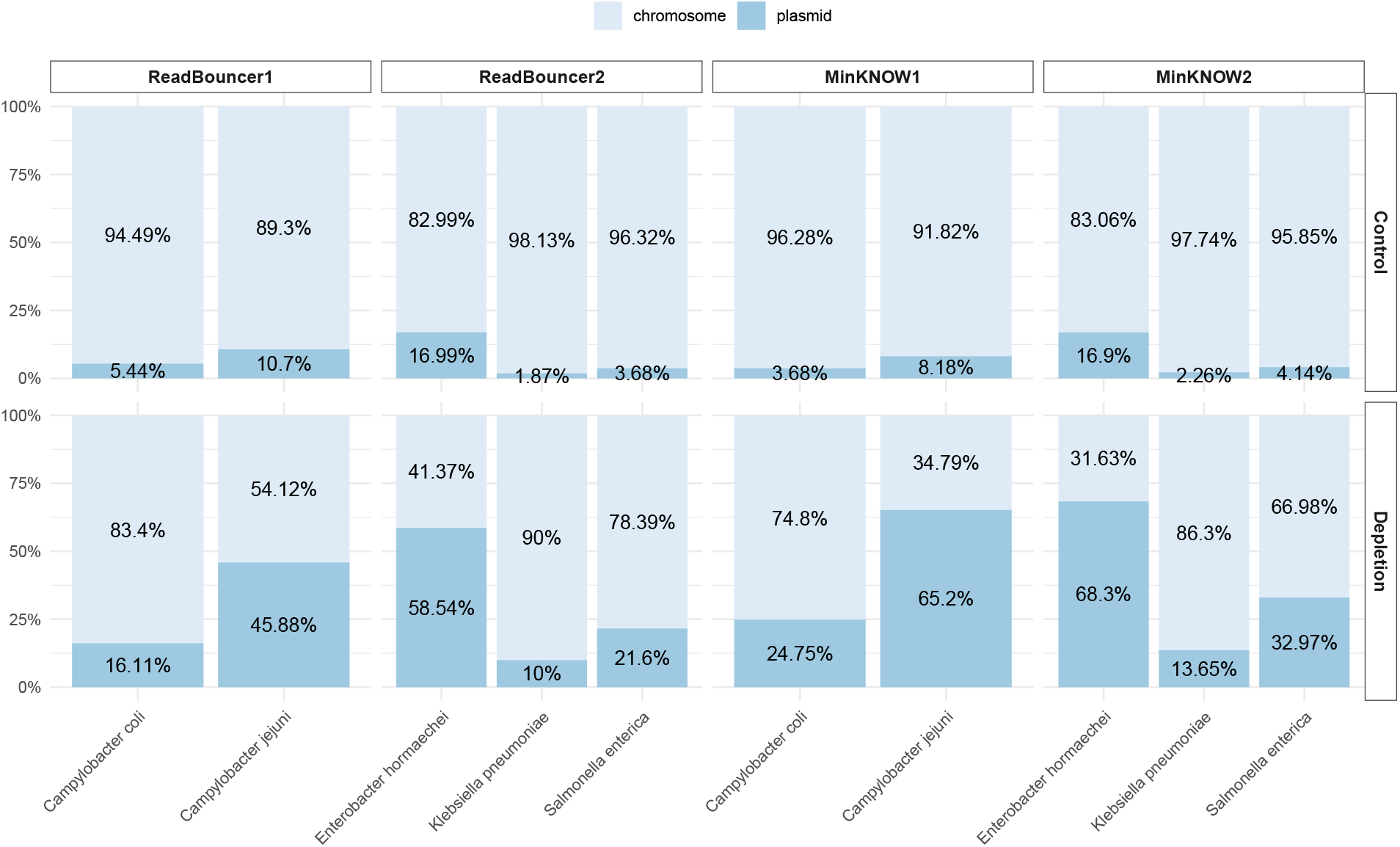
Comparison of plasmid abundances in five bacterial samples. Adaptive sampling with MinKNOW was used on flowcells MinKNOW1 and MinKNOW2 and ReadBouncer was used as adaptive sampling tool on flowcells ReadBouncer1 and ReadBouncer2. For all experiments, plasmid abundances for each sample were measured after 24 hours of sequencing for control regions and adaptive sampling regions (Depletion). Plasmid abundances are highest when using MinKNOW for depletion of chromosomal nanopore reads.

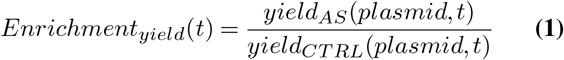

where *yield*_*AS*_(*plasmid, t*) is the number of sequenced plasmid bases of a strain from the adaptive sampling region at time point *t* and *yield*_*CT RL*_(*plasmid, t*) is the number of sequenced plasmid bases of a strain from the control region (without adaptive sampling) at time point *t*.

Similarly, we calculated the enrichment by the number of reads,

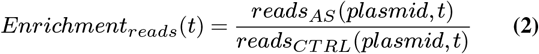

and the enrichment by the mean depth of coverage of the plasmid reference sequences.

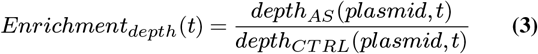

According to the definitions above, *reads*_*AS*_(*plasmid, t*) and *reads*_*CT RL*_(*plasmid, t*) represent the number of reads from the adaptive sampling (AS) or control region (CTRL) that map to a plasmid of a given bacterial strain at experiment time point *t*. Further, *depth*_*AS*_(*plasmid, t*) denotes the mean sequencing depth of plasmids from a strain using mapping data from the adaptive sampling region at time point *t* and *depth*_*CT RL*_(*plasmid, t*) is the mean sequencing depth of plasmids on the control region at time point *t*. Here, we used `samtools coverage` (27) to calculate the mean depth of coverage of every species’ plasmid reference at each time point for the control and adaptive sampling regions. The different enrichment factor values calculated for each bacterial sample at any of the 48 time points were plotted and shown in Figure 8.

For the plots of active channels over time (Figure 7), a channel was defined as active from the beginning of the experiment, up until the time it sequenced its final molecule (as long as it sequenced at least one molecule). The enrichment by composition shown in Figure 6 (a) and (b) was calculated by dividing the relative plasmid abundance from adaptive sampling regions by the relative plasmid abundance from control regions, both shown in Figure 5. We compared observed enrichment by composition and yield against predicted enrichment values using the mathematical model from Martin et al.. To calculate predicted enrichment values, we used the recommended sequencing speed of 420 bp/sec, capture time of 0.5 seconds, decision time of 1 second, and mean read lengths for each bacterial sample as provided in Table 1, and plasmid abundances of control regions for each sample as shown in Figure 5.

**Fig. 6.**
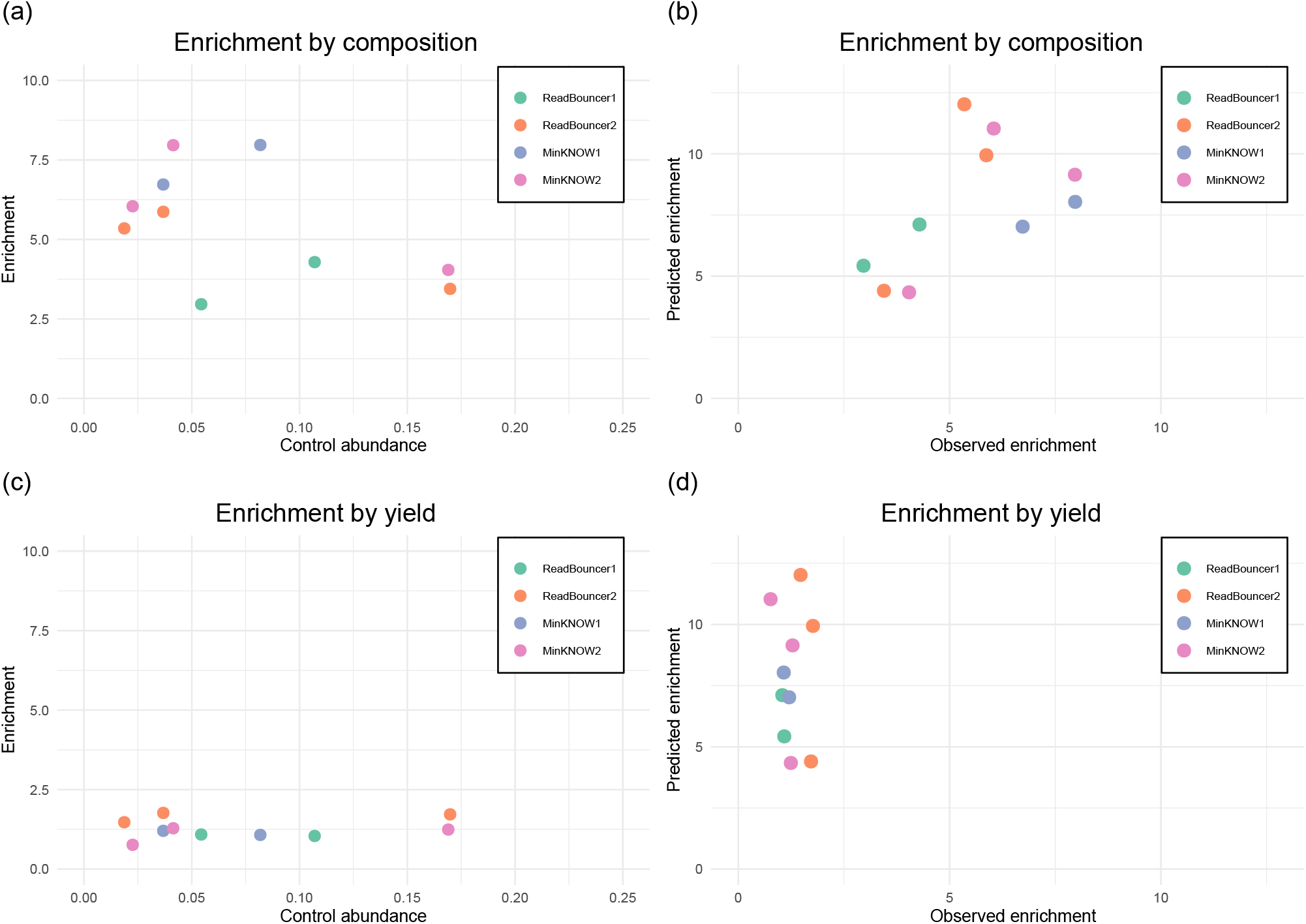
Scatterplots for plasmid enrichment by composition and yield. (a) Enrichment factor by composition against relative abundance. Each point represents a bacterial sample, with the position on the x-axis indicating the original relative abundance of plasmids in the sample and the position on the y-axis indicating the enrichment factor obtained. (b) Correlation between observed enrichment values by composition and predicted enrichment values by the mathematical model (Pearson’s r of 0.55). (c) Enrichment factor by yield against relative abundance. Each point represents a bacterial sample, with the position on the x-axis indicating the original relative abundance of plasmids in the sample and the position on the y-axis indicating the enrichment factor obtained. (d) Correlation between observed enrichment values by yield and predicted enrichment values by the mathematical model (Pearson’s r of -0.07).

**Fig. 7.**
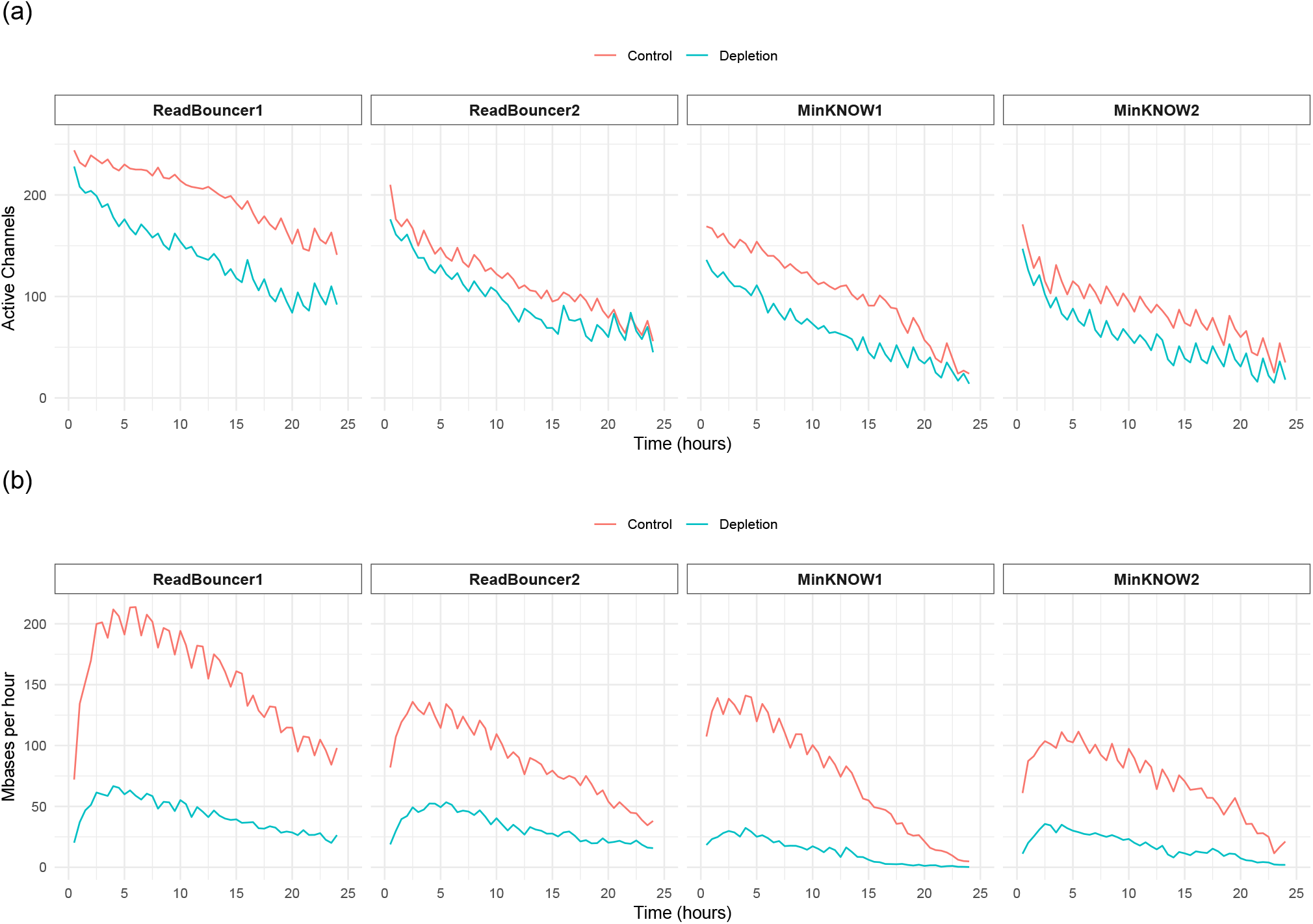
Comparison of active channels and yield between control and adaptive sampling (depletion) regions in all four experiments. **(a)** Plots showing how the number of active channels varies with time in adaptive sampling (depletion) and control regions. There are more active sequencing channels in control regions on all four flow cells **(b)** Hourly yields from depleted channels vs control channels. Usage of adaptive sampling results in lower overall sequencing yield compared to normal sequencing.

**Fig. 8.**
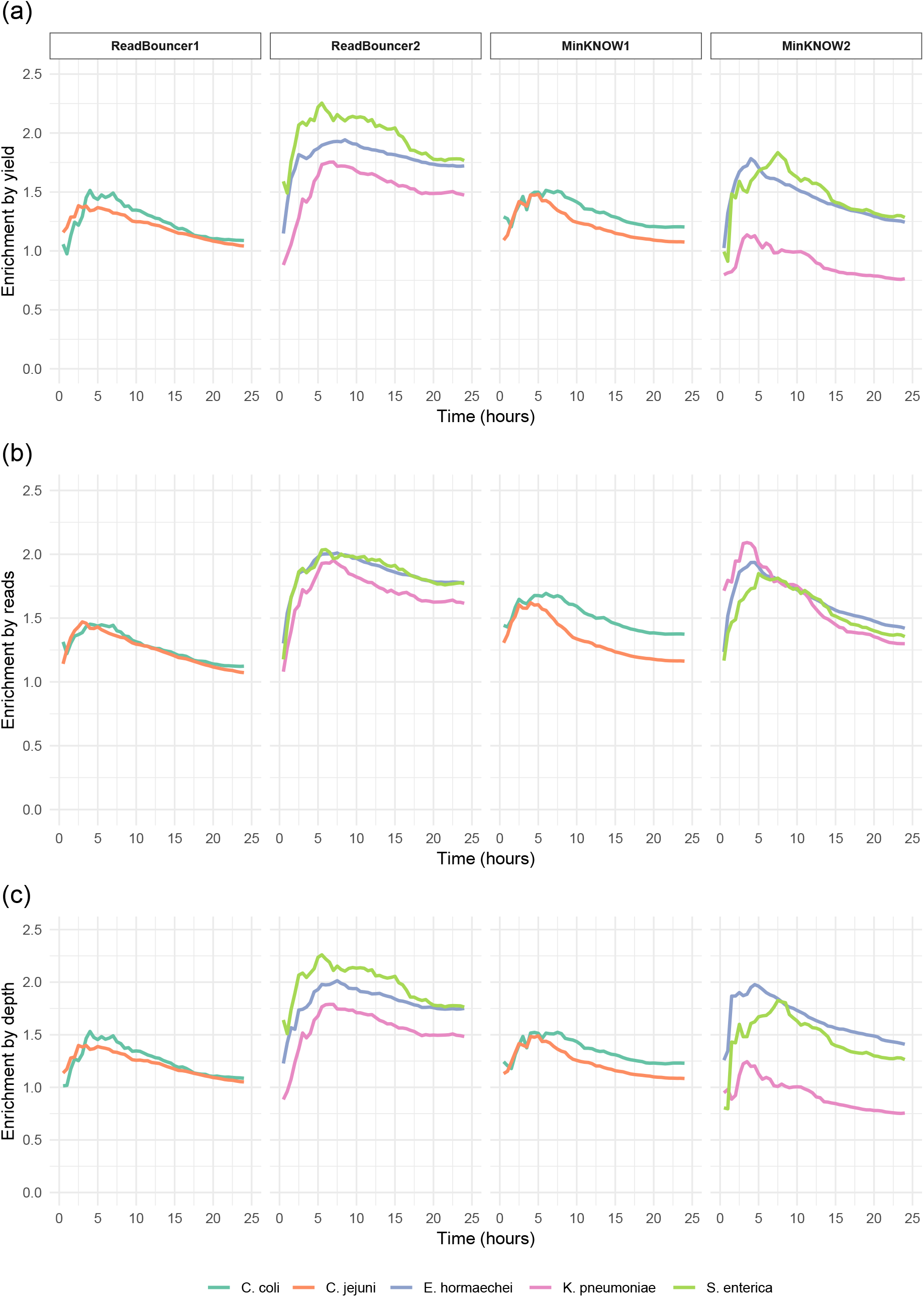
Comparison of enrichment in five bacterial samples. **(a)** Enrichment by number of sequenced plasmid bases for all five bacterial strains across the four sequencing runs. **(b)** Enrichment by number of plasmid reads for the five bacterial strains across all four sequencing runs. **(c)** Enrichment by mean depth of coverage of plasmid references for the five bacterial strains across the four sequencing runs. All strains but the *Klebsiella pneumoniae* sample, where MinKNOW was used for adaptive sampling, show a slight enrichment.

Since we expect plasmid sequences in our use case scenario to be usually unknown, we also did a *de novo* assembly of the demultiplexed fastq files, containing all reads sequenced after one and two hours of sequencing. This helps us to estimate the time required to obtain high-quality plasmid assemblies. Therefore, we assembled all demultiplexed nanopore reads from control and adaptive sampling regions separately using `Flye/metaFlye` assembler (v2.9.2, parameter “–meta”) (28, 29). Then, we polished the obtained `metaFlye` assemblies with one round of `Medaka consensus` (v.1.8.0, default parameters, model r941_min_sup_g507, Oxford Nanopore technologies) using the same nanopore read set. We assessed the quality of the final assemblies with `Quast` (v5.2.0) (30) and combined reported metrics like mean depth of coverage for both time points.

## Results

### Reduced sequencing yield but same data quality with expired flow cells

In this study, we present the application of nanopore adaptive sampling on the *in-silico* enrichment of plasmids by depleting chromosomal reads during the sequencing of bacterial isolates. Therefore, we separately sequenced five bacterial strains *Campylobacter jejuni, Campylobacter coli, Salmonella enterica, Enterobacter hormaechei* and *Klebsiella pneumoniae* on four different flow cells, each divided into an adaptive sampling and a control region. We will refer to the flow cells according to the adaptive sampling tool used. This section will provide an overview of the general sequencing run and sample metrics of the control regions to assess the quality of the four sequencing runs.

First, we investigated the number of active pores on each flow cell (Figure 1 c) and the sequencing yield from the control regions in terms of the number of base pairs and the number of reads (Figure 2 (a,c)). We see that flow cell ReadBouncer1 has the highest number of active sequencing pores (1,153) at the start of the run, while the other three flow cells have between 557 and 718 active pores. The fewer active pores can be explained using expired flow cells for these three sequencing runs. Consistent with the number of active pores, flow cell ReadBouncer1 has the highest overall sequencing yield (about 4.5 Gigabases). Surprisingly, flow cell MinKNOW2 results in significantly less yield than ReadBouncer2 (2 Gigabases vs. 3 Gigabases), although having a higher number of active pores (718 vs. 636) at the start of the sequencing run. This shows that the number of active pores does not necessarily correlate with flow cell yield for expired flow cells. The yield on control regions is much higher for all flow cells than on adaptive sampling regions. This observation is consistent with previous studies (17, 21) and can be explained by more overall time spent capturing a new molecule after rejecting one from a pore. In this context, we also see on all four flow cells a higher number of reads sequenced on the adaptive sampling regions than on the control regions (Figure 2 (c)). Thus, many reads are classified as chromosomal by the adaptive sampling tools and rejected from the pores, leading to more reads sequenced on the adaptive sampling regions. Here, the flow cell run ReadBouncer2 has a higher number of reads on the adaptive sampling region than ReadBouncer1. This results from a larger number of chromosomal reads that were rejected on the adaptive sampling region of flow cell ReadBouncer2 (approx. 370,000) than on the adaptive sampling region of ReadBouncer1 (approx. 350,000).

To further assess and compare the quality of the four sequencing runs, we look at the read length and quality from the control regions of the sequencing runs. The contour plots in Figure 3 show that for all four sequencing runs, a large proportion of reads has a mean Phred quality between 12 and 15. Only for runs ReadBouncer1 and MinKNOW2, we observe a significant proportion of reads with low mean phred quality between 5 and 7. This suggests no drop in per-read quality when using expired flow cells for normal sequencing. We can also not observe any read length-related quality drop for the expired flow cells. Regarding read lengths, we can only compare ReadBouncer1 with MinKNOW1 and ReadBouncer2 with MinKNOW2. For ReadBouncer1 and MinKNOW1, both having the same *Campylobacter* samples sequenced, we see a larger proportion of longer reads above 10,000 base pairs for ReadBouncer1. In order to investigate that difference, we looked at per-sample read length metrics, which are provided in Table 1. First, the sequencing data after base calling from the control regions show a large difference in mean and median read lengths and the standard deviation for the two *Campylobacter* samples. Here, the application of the MagAttract HMW Genomic Extraction Kit (Qiagen) results in larger read lengths for the *Campylobacter jejuni* samples. We also see a difference between the read lengths of the same sample from the two *Campylobacter* sequencing runs (ReadBouncer1 and MinKNOW1), particularly for *Campylobacter coli*. However, there is no trend that read lengths on expired flow cells are generally shorter because the read lengths for *Campylobacter coli* are longer on the expired flow cell MinKNOW1 when compared to ReadBouncer1. This suggests that sample handling and preparation, as well as the used barcodes, could have more influence on the read length than flow cell expiration. For the other two flow cells, we observe that the mean read quality is better for ReadBouncer2, and the read lengths are longer for each sample when compared to MinKNOW2. Although both flow cells were expired, we cannot say whether the flow cells’ quality could have influenced both metrics. However, the number of active pores for MinKNOW2 at the sequencing start was higher than that for ReadBouncer2, which suggests that this number is not a reliable indicator of the quality of the sequenced data.

### Adaptive sampling reduces the number of active channels and sequencing yield, but not read quality

One of the major aspects of our study is the investigation of the impact adaptive sampling has on expired nanopore flow cells. In Figure 2, we see that for all four flow cells, the sequencing yield on adaptive sampling regions is significantly reduced in comparison to control regions. This observation aligns with previous studies (17, 21) and originates from a reduced overall time spent for sequencing the DNA and more overall time needed to capture the DNA molecules when adaptive sampling is applied. However, we do not see a higher relative yield reduction for adaptive sampling regions on expired flow cells ReadBouncer1, MinKNOW1, and MinKNOW2.

In order to check if adaptive sampling leads to faster pore exhaustion on expired flow cells, we further investigated the effect of adaptive sampling on the number of active sequencing channels and yield in sequenced Mbases per hour (see Figure 7). Comparing the four experiments, we consistently observe fewer active sequencing channels on flowcell regions with adaptive sampling than in control regions across all experiments. In summary, we find between 1.4 to 2.6 times more active channels in control regions than in adaptive sampling regions. However, we could not detect bigger systematic differences in active sequencing channels on expired flow cells when compared to the fresh flow cell ReadBouncer1. Finally, we also compared the average read quality scores from reads sequenced on control regions (Figure 3) with those sequenced on adaptive sampling regions (Figure 4). This comparison shows no significant loss in read quality when applying adaptive sampling to expired flow cells.

### Rejecting chromosomal reads increases the relative plasmid abundance

In our four experiments, we investigate the potential enrichment of plasmid sequences in bacterial samples by rejecting the chromosomal reads using adaptive sampling. First, we calculated the percentage of sequenced chromosomal and plasmid base pairs for each sample from the adaptive sampling and control regions. We refer to the percentage of plasmid base pairs as the relative plasmid abundance in a sample. In Figure 5, we see that after 24 hours of sequencing, adaptive sampling increases the relative abundance of plasmid base pairs (bp) for all samples on the four flow cells. For instance, we could increase the abundance of *Campylobacter coli* plasmid bases from 3.68% to 24.75% when rejecting chromosomal reads with `MinKNOW`. We further observe that plasmid abundances are much higher when using `MinKNOW` instead of `ReadBouncer` for adaptive sampling.

We further examined whether the plasmid enrichment by composition and yield we observe in our experiments corresponds to the predicted enrichment by the mathematical model proposed by Martin et al.. We calculated the enrichment by composition by dividing the relative plasmid abundance from adaptive sampling regions by the relative plasmid abundance from control regions. Accordingly, we calculated the enrichment by yield using the number of sequenced plasmid base pairs from adaptive sampling and control regions. As predicted by the model, the enrichment factor was higher for samples with less abundant plasmids (Figure 6 (a)). The highest levels of enrichment by composition were obtained using `MinKNOW`, which can be explained by faster rejection decisions. The predictions from the mathematical model by Martin et al. correlated moderately with our observations (Pearson’s *r* = 0.55) as shown in Figure 6 (b). In contrast, the original plasmid abundance has no impact on the enrichment by yield (Figure 6), with enrichment by yield being significantly less than enrichment by composition. We also noticed that the predicted enrichment values by the model do not correlate with the observed enrichment values by yield (Pearson’s *r* = −0.07, Figure 6 (d)).

### Effective enrichment of plasmids by yield, read number and mean depth of coverage

We examine the effective plasmid enrichment at different time points of sequencing for each experiment by calculating the plasmid enrichment for the five bacterial species in 30-minute intervals. According to equation 1, the enrichment by yield is the ratio of cumulative plasmid bases from the adaptive sampling region and the control region at time point *t*. We calculate the enrichment by the number of plasmid reads and mean depth of coverage in the same manner as proposed by equations 2 and 3. Figure 8 (b) illustrates that we obtain an enrichment of plasmid reads for all samples in all experiments at any given time point. This observation confirms that the number of sequenced plasmid reads can be increased by using adaptive sampling. We can see the same effect for the enrichment by yield (Figure 8 (a)) for all but one sample. For the *Klebsiella pneumoniae* sample of flow cell MinKNOW2, we observe that the number of plasmid bases from adaptive sampling is less than that from the control channels. Thus, we failed to obtain an enrichment of *Klebsiella pneumoniae* plasmids in that experiment where we used `MinKNOW` to deplete chromosomal reads. For all other samples, we observe an enrichment of 1.1x to 1.8x after 24 hours, corresponding to 10*−*80% more plasmid data when using adaptive sampling, even when using expired flow cells with reduced active pores.

We further investigated the difference in enrichment between the same samples from experiments ReadBouncer2 and MinKNOW2. First, flow cell MinNKOW2 has fewer active sequencing channels and produces less sequencing yield than the flow cell from experiment ReadBouncer2 (see Figure 7). Figure 4 also illustrates that the average read quality in the adaptive sampling region of flow cell MinKNOW2 is smaller than for flow cell ReadBouncer2. Both observations suggest a decreased pore quality of flow cell MinKNOW2. Although this might explain the reduced enrichment by yield in this experiment, it does not explain why there is an effective depletion of plasmid bases for *Klebsiella pneumoniae* in experiment MinKNOW2. Thus, we identified all reads from the final output that mapped against the *Klebsiella pneumoniae* plasmids but were rejected by `MinKNOW`. We extracted these falsely rejected plasmid reads of *Klebsiella pneumoniae* and mapped them to the corresponding bacterial chromosome reference sequences with `minimap2`. Using `samtools depth`, we could identify four regions (between 829 and 2,101 bp long) on the *Klebsiella pneumoniae* chromosome with read depth*≥*10. These findings reveal regions of high identity between *Klebsiella pneumoniae* plasmid targets and non-target chromosome sequences. Such similar regions between target and non-target sequences pose a challenge for the application of nanopore adaptive sampling and potentially lead to an increased number of falsely rejected target reads. Here, it seems that ReadBouncer can avoid a high number of false rejections by using longer read prefixes (see Figures **??** and 4) for making rejection decisions. Our observations suggest that using more sequence information by increasing the chunk size for adaptive sampling with MinKNOW could circumvent such issues.

### Adaptive sampling helps improving plasmid assemblies

Our experiments demonstrated an effective enrichment of plasmids after 2-5 hours by using adaptive sampling. Since plasmid assemblies are possible after 3-4 hours of sequencing without adaptive sampling (10), we wanted to see if adaptive sampling enables faster plasmid assemblies. In order to evaluate the effect of adaptive sampling on the assembly of low-abundant plasmids, we took the reads available after one hour and two hours of sequencing from the adaptive sampling and control regions two of the bacterial isolates, *Salmonella enterica* and *Enterobacter hormaechei*. We did not include the *Campylobacter* strains in this analysis because the coverage of plasmids from sequencing only two isolates on a fresh flow cell (sequencing run ReadBouncer1) was extraordinarily high and assembly statistics would not be comparable between the sequencing runs ReadBouncer1 and MinKNOW1. We also did not include *Klebsiella pneumoniae* because of the findings mentioned in the last subsection that could bias our analysis.

We separately assembled all reads for the control and adaptive sampling regions using `metaFlye` assembler (29). After one round of polishing with `Medaka consensus`, we measured quality metrics for the final assemblies using `Quast` (30). Table 2 shows results after one and two hours of sequencing the two bacterial isolates from sequencing runs ReadBouncer2 and MinKNOW2. In all cases, plasmid assemblies were improved by using adaptive sampling, with reduced numbers of mismatches and indels for plasmid assemblies from the adaptive sampling regions.

**Table 2.** Plasmid Assembly statistics of adaptive sampling and control region after one and two hours of sequencing for two different bacterial isolates from sequencing runs ReadBouncer2 and MinKNOW2. All reads from the adaptive sampling and control regions were separately assembled for *Salmonella enterica* and *Enterobacter hormaechei* using `Flye` assembler. Assembly statistics provided by `Quast` show better results for plasmid assemblies from adaptive sampling than control regions.

|  | *Salmonella enterica* |  |  |  | *Enterobacter hormaechei* |  |  |  |
| --- | --- | --- | --- | --- | --- | --- | --- | --- |
|  | ReadBouncer |  | MinKNOW |  | ReadBouncer |  | MinKNOW |  |
| Adaptive Sampling (channels 1-256) |  |  |  |  |  |  |  |  |
| Time (hours) | 1 | 2 | 1 | 2 | 1 | 2 | 1 | 2 |
| Ref. plasmids | 1 | 1 | 1 | 1 | 4 | 4 | 4 | 4 |
| Assembled plasmid contigs | 1 | 1 | 2 | 1 | 4 | 4 | 6 | 5 |
| Plasmid reads | 108 | 277 | 80 | 194 | 1,383 | 3,656 | 1,469 | 3,831 |
| Ref. avg. coverage depth | 11 | 29 | 5 | 19 | 14 | 43 | 13 | 37 |
| Ref. coverage $\geq 5x(\%)$ | 100 | 100 | 84.67 | 100 | 98.41 | 99.84 | 91.99 | 99.84 |
| Ref. coverage $\geq 10x(\%)$ | 70.81 | 100 | 2.8 | 100 | 54.28 | 99.84 | 53.12 | 97.68 |
| Mismatches per 100kb | 136 | 126 | 358 | 132 | 55 | 26 | 94 | 38 |
| Indels per 100kb | 653 | 652 | 895 | 656 | 89 | 25 | 136 | 40 |
| Control (channels 257-512) |  |  |  |  |  |  |  |  |
| Time (hours) | 1 | 2 | 1 | 2 | 1 | 2 | 1 | 2 |
| Ref. plasmids | 1 | 1 | 1 | 1 | 4 | 4 | 4 | 4 |
| Assembled plasmid contigs | 1 | 1 | 1 | 1 | 6 | 4 | 6 | 5 |
| Plasmid reads | 79 | 160 | 59 | 128 | 945 | 2,136 | 971 | 2,186 |
| Ref. avg. coverage depth | 7 | 15 | 7 | 13 | 11 | 25 | 10 | 24 |
| Ref. coverage $\geq 5x(\%)$ | 98.41 | 100 | 88.38 | 100 | 82.66 | 99.34 | 73.8 | 99.84 |
| Ref. coverage $\geq 10x(\%)$ | 31.21 | 96.98 | 21.07 | 87.24 | 25.46 | 89.4 | 23.06 | 90.7 |
| Mismatches per 100kb | 219 | 133 | 397 | 144 | 193 | 31 | 306 | 93 |
| Indels per 100kb | 744 | 654 | 764 | 652 | 238 | 43 | 488 | 137 |

This can be explained by an increased sequencing depth and reference coverage due to the plasmid enrichment by adaptive sampling. However, we recognize some gaps in *Enterobacter hormaechei* plasmids assembled using reads from the adaptive sampling region of run MinKNOW2. This can be caused by similar regions between target and non-target sequences, as we have observed for *Klebsiella pneumoniae*. In general, adaptive sampling can improve plasmid assemblies and enables assemblies even after 2 hours of sequencing on flow cells with fewer active pores.

## Discussion

Recent studies have demonstrated the utility of adaptive sampling for the enrichment of underrepresented sequences in various applications, such as host depletion in human vaginal samples or antibiotic resistance gene enrichment in metagenomics samples. In this study, we examine the potential of adaptive sampling for the enrichment of low-abundant plasmid sequences by rejecting chromosomal sequences in bacterial isolate samples. We demonstrate the possibility of using even older or expired flow cells with fewer active sequencing pores for the *in-silico* enrichment via adaptive sampling. Since we wanted to know if enrichment is independent of the adaptive sampling tool, we evaluated plasmid enrichment for two tools, namely `ReadBouncer` and ONT’s `MinKNOW` sequencing control software. Although we observed different levels of plasmid enrichment, the tools consistently enriched for low abundant plasmid sequences. Our study was by no means designed to benchmark different adaptive sampling tools, which would require the inclusion of more tools and a setup that ensures that all tools use the same amount of sequence information for making rejection decisions.

The enrichment by yield, the most critical value for researchers, lies for all but one sample in our experiments between 1.1x and 1.8x after 24 hours of sequencing on an ONT MinION sequencing device. We also demonstrated that the difference between enrichment by yield, number of reads and mean depth of coverage is negligible in all our samples. High-quality assemblies of plasmids are possible within two hours of sequencing with adaptive sampling and show even better results than plasmid assemblies without adaptive sampling. These results reflect the benefit of adaptive sampling in assembling low-abundant plasmid sequences. Since we sequenced three bacterial isolates on only half a reused flow cell, we reason that up to 20 bacterial isolates can be sequenced on a flow cell with adaptive sampling for plasmid enrichment.

Our experiments showed that expired flow cells with a decreased number of active pores could be used in combination with adaptive sampling. Previous studies demonstrated that the number of active sequencing pores decreases faster when using adaptive sampling. Although we show the same trend in our study, we do not see a negative impact on the enrichment of target sequences and the average quality of sequenced reads. Thus, we encourage researchers to use flow cells with reduced active pores in adaptive sampling experiments for more sustainable lab experiments and cost savings in core facilities and larger research institutions.

Our results show that rejecting chromosomal sequences with adaptive sampling increases the abundance of plasmid sequences in the final output. Depending on the plasmid abundance in the original sample, the values for plasmid enrichment by composition are between 2.5x and 8x. These observations moderately correlate with the predictions from the mathematical model proposed by Martin et al.. Furthermore, a consistent enrichment of plasmid sequences with regard to the number of base pairs, number of reads, and depth of coverage was shown by using adaptive sampling. Independent of the size of the sequencing libraries, we could increase the amount of sequenced plasmid base pairs by 10-80% after 24 hours of sequencing. However, in one experiment, we recognized the depletion of plasmid sequences of *Klebsiella pneumoniae* after 24 hours when ONT’s `MinKNOW` was used as an adaptive sampling tool. Our investigations reveal that regions with high sequence identity located both on the chromosome and the plasmid lead to false read rejections, which result in a depletion of the targeted plasmid sequences. This highlights potential issues with the usage of nanopore adaptive sampling and sounds a note of caution if target and non-target sequences are similar. We hypothesize from our findings that using larger read chunks for making rejection decisions could circumvent this issue. However, such an examination is beyond the scope of this study and needs systematic investigations to find the optimal read chunk length that minimizes false rejections decisions while still rejecting unwanted reads fast enough to obtain sufficient enrichment.

Both adaptive sampling tools used in this study need known reference sequences to reject the chromosomal reads. If the bacterial species in the given sample are unknown, a more extensive reference database of all potential bacterial chromosome references must be used to enrich plasmids successfully. Alternatively, researchers could also do a targeted enrichment of the plasmids by using plasmid databases such as PLSDB (31) and reject all reads that do not match the database. However, this approach risks missing unknown plasmids not covered by the database. Using specific plasmid markers, like the origin of replication, for correctly classifying unknown plasmids is also tricky in an adaptive sampling experiment. Here, the specific markers would need to be located on the first 1000 bp of the read to prevent false rejection of plasmid reads. These limitations reinforce the need for improved classification algorithms that can even classify reads from unknown plasmids based on the raw nanopore signals.

We envisage several applications for the *in-silico* enrichment of plasmids in the near future. One possibility is the surveillance of plasmid outbreaks in hospital settings. Here, clinicians are interested in studying the transmission of specific antibiotic resistance genes harboring plasmids from one bacterial species to another. Such community transmissions can indicate the selection pressure on bacteria caused by antibiotic pharmaceuticals and help decide on the corresponding drugs’ future usage.

Another possible application of adaptive sampling is the improvement of known bacterial assemblies. In this study, we demonstrated the improved time-to-assembly of plasmids by depleting the known bacterial chromosomes. We plan to develop a pipeline for the real-time *de novo* assembly of bacterial isolates in the future. Using adaptive sampling, we could reject reads that cover assembled regions with a minimum depth of coverage, enriching for unseen or assembled regions with low sequencing depth. In such a way, we could complement the dynamic re-sequencing framework BOSS-RUNS (32) with a dynamic *de novo* adaptive sampling framework. We believe this could improve both the quality of bacterial and plasmid assemblies as well as metagenomics assemblies of unknown bacterial species.

## Conclusions

Using nanopore adaptive sampling, we demonstrated an *in-silico* enrichment of low abundant plasmid sequences in known bacterial isolates by rejecting chromosomal sequences. We discovered that enrichment could also be achieved with expired flow cells, which, combined with adaptive sampling, shows the potential for cost savings in clinical and laboratory sequencing of bacterial pathogens. The combination of flow cell re-usage and omittance of time-consuming wet lab enrichment of plasmids make the application of plasmid sequencing more attractive for routine clinical diagnostics.

We also observed that similar regions between target and non-target sequences pose a challenge for adaptive sampling tools that use mapping algorithms for making rejection decisions. In such scenarios, adaptive sampling could even deplete target sequences if similar regions are not masked in the non-target reference sequences. By performing targeted enrichment of low abundant plasmids, we significantly improved the quality of *de novo* assemblies of the plasmids and reduced the sequencing time needed to achieve this quality. We conclude that nanopore adaptive sampling will be useful for many bacterial plasmid sequencing studies.

## Availability of data and materials

Scripts and software used in the analysis are available in the GitHub repository https://github.com/JensUweUlrich/PlasmidEnrichmentScripts, as described in the methods.

The sequence datasets generated and analyzed during the current study are available in the NCBI BioProject repository under accession number PRJNA862336.

## ACKNOWLEDGEMENTS

The authors thank Martin Hölzer, Matthew Huska, and Aleksandar Radonic (RKI) for valuable discussions and comments on the usage of nanopore adaptive sampling. We thank the Genome Sequencing Unit at Robert Koch Institute for sequencing the bacterial strains.

